# Genomic insights into sex-linked regions and dioecy in the ivory palm (*Phytelephas aequatorialis*)

**DOI:** 10.64898/2026.09.17.752333

**Authors:** Perla Farhat, Julie Orjuela, Hugo Tessarotto, Jos Käfer, Thierry Beulé, Galilea Orellana-Vera, Romain Guyot, Rommel Montúfar, Frederique Aberlenc

## Abstract

**Background and Aims:** Dioecy has evolved independently multiple times in Palms, yet the genetic mechanisms underlying sex determination remain poorly understood across the family. Current knowledge is limited to genera from the subfamily Coryphoideae, in which *Phoenix* and *Kerriodoxa* independently recruited the same genomic region as the sex determining locus, suggesting a potentially convergent evolutionary pathway. Whether this pattern extends across the family remains unknown. Here, we investigate the genetic basis of sex determination in *Phytelephas aequatorialis*, from the Ceroxyloideaea palm subfamily, to identify its sex chromosome system, sex-linked genomic region, and candidate sex-determining genes.

**Methods:** Using exon capture sequencing data from male and female individuals of *P. aequatorialis*, we analyzed genome wide sequence polymorphisms to detect signatures of sex linkage.

**Key Results:** Our analyses reveal an XY sex chromosome system in *Phytelephas*, similar to that reported in the few dioecious palm species investigated to date. However, sex linkage maps to a distinct genomic region, indicating an independent origin of the sex-determining locus. We further identified 16 high confidence sex-linked genes with key roles in developmental processes and hormone signaling pathways, making them key candidates for involvement in sex determination. Sex-specific genetic markers were successfully developed from a subset of these genes.

**Conclusions:** Our study suggests that in *P. aequatorialis*, the independent evolution of sex separation has relied on distinct genomic regions and genes, highlighting multiple evolutionary pathways leading to dioecy across the palm family. Comparative genomic analyses across additional lineages will be essential to reconstruct the evolutionary history of sex chromosomes and sex separation in this family.

## Introduction

Sexual systems in angiosperms vary in the spatial distribution of male and female reproductive functions, ranging from bisexual flowers to different forms of unisexuality (Barrett 2002). Most species are hermaphroditic, producing flowers that contain both male and female organs, whereas unisexual systems involve the separation of sexes either within individuals (monoecy) or among individuals (dioecy). Dioecy is relatively rare, occurring in approximately 6% of angiosperm species, yet it is phylogenetically widespread, having evolved independently across various lineages (Renner and Ricklefs 1995). It has been reported in at least 157 families and encompassing around 37% of the angiosperm diversity at the family level (Renner and Ricklefs 1995; Hultine *et al*. 2016; Sauquet *et al*. 2017; Marais *et al*. 2025). The transition to dioecy is often associated with potential several evolutionary advantages, including enhanced outcrossing, increased genetic diversity, reduced inbreeding depression, and more efficient allocation of reproductive resources (Barrett *et al*. 2010; Pannell and Jordan 2022; Osterman *et al*. 2024).

Dioecy is thought to have evolved repeatedly and relatively recently from hermaphroditic ancestors (Sauquet *et al*. 2017; Marais *et al*. 2025). These independent transitions to dioecy across angiosperms therefore raise the question of whether they rely on shared genetic mechanisms or instead involve distinct molecular pathways recruited in different lineages (Ming *et al*. 2011; Charlesworth 2016; Muyle *et al*. 2017). Recent advances in genomics, particularly the development of high-throughput sequencing technologies and long-read approaches enabling chromosome-scale genome assemblies, have substantially improved our ability to investigate the genetic basis of sex determination (Hobza *et al*. 2024; Kudoh *et al*. 2026). These technological developments are driving a shift from traditional genetic and cytological approaches toward integrative, genome-wide analyses, including the characterization of structural variation, recombination landscapes, and epigenomic regulation (Hobza *et al*. 2024). However, despite these technological advances, our understanding of sex determination mechanisms in plants remains limited. To date, sex determination mechanisms have been characterized in fewer than 100 angiosperm species, representing only a small fraction of the estimated dioecious species in this group (ca. 15 000) (Charlesworth 2016). Among the studied species, the majority exhibit an XY system with male heterogamety (Ming *et al*. 2007; Charlesworth 2016). ZW sex chromosome systems (female heterogamety) have also been reported in some lineages, yet the current data are insufficient to assess their overall prevalence (Muyle *et al*. 2017; Käfer *et al*. 2022; Renner 2025).

In studied species with heteromorphic sex chromosomes (Y or W chromosomes), a non-recombining region, referred as the sex-determining region (SDR), has been characterized (Ming *et al*. 2007; Charlesworth 2016). The establishment of recombination suppression within the SDR has been investigated in only a limited number of systems and is often associated with structural genomic changes, such as inversions and sequence expansions (Bergero and Charlesworth 2009; Bačovský *et al*. 2020). Moreover, SDR holds candidate sex-determining genes which have been identified in a few dioecious species, mainly involving pathways related to hormonal regulation (e.g. cytokinin signaling) and floral organ development (Harkess *et al*. 2017; Wybouw and De Rybel 2019; Varkonyi-Gasic *et al*. 2021). However, the multiple and independent origins of dioecy across angiosperms highlight the need for broader comparative studies across phylogenetically diverse lineages to fully elucidate the mechanisms underlying the evolution of unisexuality.

The palm family (Arecaceae), comprising approximately 2 600 species, exhibits remarkable diversity in sexual systems, with ca. 52% monoecious, ca. 30% dioecious, and ca. 17% hermaphroditic species (Dransfield *et al*. 2008; Nadot *et al*. 2016). Given that a majority of palm species display some degree of sex separation, this family represents a valuable model for investigating the evolution and genetic control of dioecy. Phylogenetic analyses suggest that dioecy has evolved independently multiple times within palms, arising from monoecious or hermaphroditic ancestors (Nadot *et al*. 2016). However, our current understanding of sex determination in this family remains largely restricted to one subfamily among the five present in palms, the Coryphoideae (Daher *et al*. 2010; Cherif *et al*. 2013; Torres *et al*. 2018; Tessarotto *et al*. 2025). Within this subfamily, the date palm (*Phoenix dactylifera*) has been extensively studied (Daher *et al*. 2010; Cherif *et al*. 2013; Torres *et al*. 2018; Tessarotto *et al*. 2025). Morphological analyses indicate that unisexuality results from the developmental arrest of sterile organs in male and female flowers (Daher *et al*. 2010). At the genetic level, an XY sex chromosome system has been identified in *Phoenix dactylifera*, with earlier studies estimating a ∼6 Mb non-recombining region (Cherif *et al*. 2013; Torres *et al*. 2018; Hazzouri *et al*. 2019). Recent T2T genome assemblies resolved a larger ∼15.2 Mb Y-haplotype and a ∼4.7 Mb X-haplotype on chromosome 14, together with a 1.6 Mb Y-specific inversion (Celii *et al*. 2026). Phylogenetic evidence further supports a single dioecious origin within the genus *Phoenix* (Cherif *et al*. 2016). More recently, comparative analyses within the Coryphoideae between *P. dactylifera* and the distantly related dioecious palm *Kerriodoxa elegans* revealed that the same genomic region independently evolved into a sex-linked region in both lineages, suggesting a rare case of convergent evolution (Tessarotto *et al*. 2025). This region showed to encompass sex-linked genes mainly involved in floral development and hormonal regulation networks (Tessarotto *et al*. 2025).

Despite these advances, major gaps remain in our understanding of sex determination across the palm family. In particular, little is known about other subfamilies, including Ceroxyloideae, which comprise predominantly dioecious species and whose most recent common ancestor is therefore likely to have been dioecious (Dransfield *et al*. 2008; Nadot *et al*. 2016). This subfamily includes ecologically and economically important taxa such as *Phytelephas aequatorialis* (the “ivory palm”). This species is native to humid lowland forests of western Ecuador (< 1.500 masl), where it plays a key ecological role as a food resource for wildlife. It is characterized by large pinnate leaves, basal inflorescences, and pronounced sexual dimorphism, with male inflorescences forming elongated catkin-like structures (1 to 2.5 m long) and female inflorescences arranged in head-like structures (20 to 40 cm long). Each female inflorescence develops into a spherical infructescence (30 cm in diameter), comprising approximately 20 - 25 obconical fruits, with 4 to 8 seeds per fruit. The seeds of these fruits contain a hard endosperm known as “vegetable ivory” (tagua), which is of economic and cultural importance (Koziol and Pedersen 1993; Bernal 1998; Pülschen 2000; Brokamp 2015; Escobar *et al*. 2021, 2022; Montúfar *et al*. 2022; Loayza *et al*.2024).

In this study, we investigate the genetic basis of sex determination in *P. aequatorialis*. Specifically, we aim to (i) identify its sex chromosome system and associated sex-determining region, (ii) characterize candidate sex-linked genes of this species, and (iii) assess whether the genomic region implicated in dioecy in *Phoenix* and *Kerriodoxa* has also been recruited in this phylogenetically distant lineage. Additionally, given the economic relevance of this species, we seek to develop molecular markers for early sex identification, facilitating the selection of productive female individuals in cultivation.

## Material and method

### Plant sample

Leaves from 42 samples of *Phytelephas aequatorialis* Spruce belonging to 9 populations from western Ecuador were collected and dried prior to analysis (Fig. 1, Supplementary Data Table S1). Of these, 21 samples from a single population were used to implement an exon capture approach (Supplementary Data Table S1, Pop-ExC) for the detection of sex-linked single-nucleotide polymorphisms (SNPs) and genes. The remaining samples were used to validate the sex-linked genes (Supplementary Data Table S1, VSLG) identified in the exon capture analysis through Polymerase Chain Reaction (PCR), allowing differentiation between male and female individuals.

**Figure 1:**
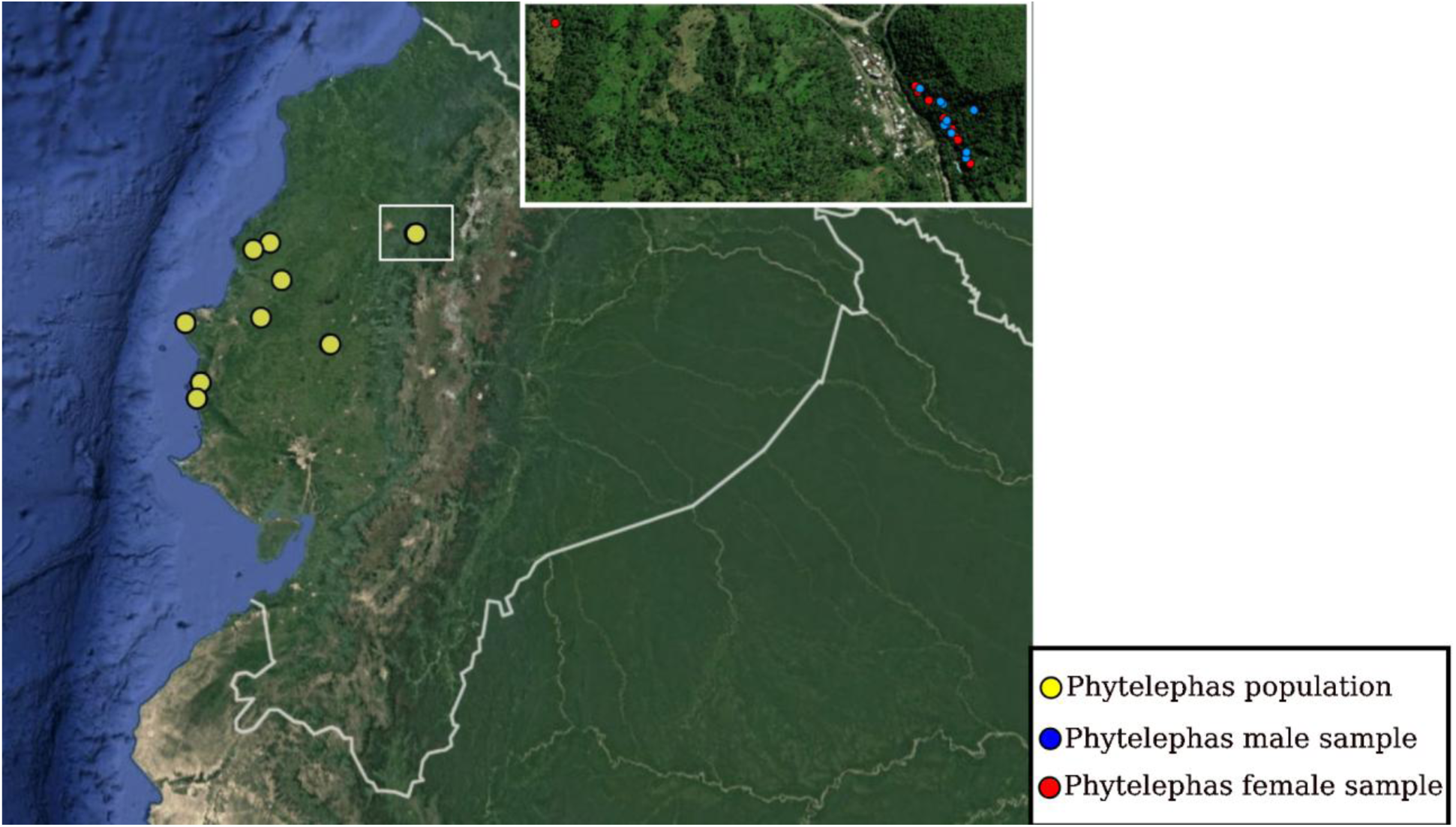
Geographical distribution of the nine *Phytelephas aequatorialis* studied populations from Ecuador. The boxed population highlights the site from which male (blue circles) and female (red circles) individuals were selected for exon capture analyses. Individuals from the remaining populations were used to validate SNP markers identified from candidate sex-linked genes.

### Exon capture targeting and sequencing

DNA was extracted from 15 mg of dried leaves of the 21 samples from Pop-ExC (11 female and 10 male individuals) with the Chemagic DNA Plant Kit (Perkin Elmer Chemagen, Germany), according to the manufacturer’s instructions. The protocol was adapted to the use of the KingFisher Flex™ (Thermo Fisher Scientific, Waltham, MA, USA) automated DNA purification workstation. We performed exon capture of *P. aequatorialis* using the PalmExon baits set, that was previously designed using the transcriptomic and genomic data of *P. dactylifera* L. cultivar ‘Khalas’ (NCBI Bioproject PRJNA83433) and was validated for its efficiency for target enrichment on other palm species (Tessarotto *et al*. 2025). The kit contains biotinylated RNA probes of 80-mers (MYcroarray, Arbor Biosciences, Ann Arbor, MI, USA) and was designed to target two exon regions of each 30 000 protein coding nuclear genes of *P. dactylifera* genome. PalmExon bait kits construction and its targeted sequences are detailed in Tessarotto et al. (2025).

For each sample, 100 ng of total DNA was fragmented using the NEBNext Ultra II FS DNA Modul (New England Biolabs, Ipswich, MA, USA), ligated to the adapters and the genomic library was constructed following published protocols (Rohland and Reich 2012; Mascher *et al*. 2013) with some modifications. All the samples from the Pop-ExC population were multiplexed into one pool and the hybridization procedure was elaborated based on the myBaits protocol v.4.01 (http://www.mycroarray.com). The final library was sequenced using the Illumina paired-end protocol (2 × 150 bp) on a HiSeq3000 sequencer (Novogene, Beijing, China).

### Data cleaning and evaluation

Raw reads were demultiplexed using Demultiplexer (https://github.com/DominikBuchner/demultiplexer). Quality control was performed both before and after read cleaning using FastQC v.0.12.1 (http://www.bioinformatics.babraham.ac.uk/projects/fastqc). Reads were trimmed to remove adapter sequences and low-quality bases using Trimmomatic v0.39 (Bolger *et al*. 2014) with the parameters SLIDINGWINDOW:4:15 and MINLEN:36. Clean reads data coverage in terms of gene number and depth was elaborated by mapping the reads on the reference genome of *P. dactylifera* ‘Barhee’ cultivar (GenBank accession: GCA_009389715.1; referred below to *P. dactylifera* BC4) using BWA-MEM2 v2.2.1 (Vasimuddin *et al*. 2019), SAMtools v1.22.1and BCFtools intersect v1.22 (Danecek *et al*. 2021).

### Variant calling and population structure analysis

Per base sequencing coverage was computed using the genomecov package of BEDTools v2.30 (Quinlan and Hall 2010). Mean coverage depth was calculated in non-overlapping 100 bp windows for each sample. In addition, the mean coverage at the gene level was estimated using only windows overlapping coding sequences (CDS) by at least 50%. The proportion of female reads per 100 bp window was calculated after correcting for differences in sequencing depth among individuals. Specifically, coverage values for each window were normalized by dividing the mean depth by the median depth across all windows for the corresponding individual.

SNPs were called using BCFtools mpileup v1.9 (Lefouili and Nam 2022). Genotypes supported by fewer than seven reads in the capture data were removed from the resulting VCF file. This filtering step primarily eliminated off-target reads, retaining variants located within the targeted CDS of *P. dactylifera*. To assess population structure and to verify that individuals did not cluster by sex, which could bias downstream analyses, we performed a principal component analysis (PCA) of genetic variation among individuals using PLINK v1.90 (Purcell *et al*. 2007). In addition, pairwise relatedness among individuals was estimated using the “--relatedness2” option in VCFtools v0.1.16 (Danecek *et al*. 2011), where values close to 0.5 indicate clones, ca. 0.25 full siblings, and ca. 0.125 second degree relatives. A heatmap of pairwise relatedness scores among the studied individuals was generated using the ggplot2 package in R (Wickham 2016).

### Determination of the sex chromosome system and sex-linked genes

Sex chromosome systems and sex-linked genes were identified using SDpop (Käfer *et al*. 2021). This method implements a population genetic framework that evaluates sex-specific genotype frequencies under alternative segregation models, including autosomal inheritance, XY or ZW gametology, hemizygosity, paralogy, and haploidy. By modeling these scenarios, SDpop provides a probabilistic assessment of sex linkage while accounting for common sources of noise in genomic data, such as genotyping errors and paralogous sequences. At the genome wide level, SDpop compares alternative sex chromosome systems using the Bayesian Information Criterion (BIC), and the system with the lowest BIC value is retained as the best-supported model for the species.

For each gene, SDpop estimates allele frequencies on the putative sex chromosome copies based on observed genotypes in females and males and computes posterior probabilities for each segregation type. Based on the retained sex chromosome systems (XY or ZW), genes showing strong support for an XY model or ZW model (posterior probability > 0.6) were retained as candidate sex-linked. For these genes we further reconstructed the consensus haplotypes (X/Y or Z/W) from allele frequency estimates. We followed the default parameters, treating alleles as fixed when their estimated frequency exceeded 0.95, assigning the major allele when frequencies ranged between 0.6 and 0.95, and marking sites with lower support as missing.

Because SDpop relies on average genotype allele frequency patterns rather than local polymorphism density, we complemented this analysis with additional sequence-based metrics. We first quantified heterozygosity at the gene level separately for males and females and calculated the relative contribution of male heterozygosity as the ratio of male heterozygosity to total heterozygosity across sexes. This statistic was examined for all genes containing at least one heterozygous site. We then estimated synonymous divergence (dS) between inferred the phased haplotypes (XY or ZW) already generated by SDpop using codeML in PAML v1.9 (Yang 2007). Sex-linked gametologs are expected to show independent evolutionary trajectories on the sex chromosomes (X/Y or ZW) following recombination suppression, leading to the accumulation of neutral substitutions and elevated dS values. In contrast, sex biased genotype frequencies caused by chance or linkage to non-recombining regions without long-term X/Y or ZW differentiation are not expected to produce substantial sequence divergence. We therefore compared the dS values of candidate sex-linked genes to the genome heterozygosity levels, using the species-specific median per gene heterozygosity as a reference. Genes with dS values exceeding the genome average heterozygosity were considered to provide stronger support for sex linkage.

To further validate the inference of sex-linked genes, we complemented the SDpop analyses and dS with iKISS (https://forge.ird.fr/diade/iKISS), which is designed to detect sex linkage from sex specific patterns of polymorphism. iKISS identifies candidate sex-linked k-mers by contrasting genotype and allele frequency distributions between female and male individuals, with a particular emphasis on sex biased heterozygosity and allele presence/absence patterns expected under XY or ZW inheritance.

Genes were classified according to combined evidence from SDpop, dS, and iKISS. SDpop posterior probabilities were divided into high (≥ 0.8), intermediate (> 0.6 and < 0.8), and low (≤ 0.6) support for sex-linked inheritance. Genes with high posterior support in SDpop (≥ 0.8), a dS value greater than or equal to the calculated threshold, and independent support from iKISS based on a high probability of sex-linked k-mers were classified as high-confidence sex-linked (HCSL). Genes with high posterior support in SDpop (≥ 0.8) were classified as probably sex-linked (PSL) when either (i) their dS value was below the threshold, regardless of iKISS classification, or (ii) their dS value was greater than or equal to the threshold but they were not identified as sex-linked by iKISS. Genes with intermediate posterior support in SDpop (> 0.6 and < 0.8), together with a dS value greater than or equal to the threshold, were also classified as PSL, regardless of whether they were supported by iKISS. In addition, genes identified exclusively by iKISS with a high probability of sex-linked k-mers, but not analyzed by SDpop, were classified as PSL. Such cases may represent highly diverged, repetitive, or structurally variable sex-linked regions that are difficult to detect using SNP-based population-genetic models but remain identifiable through sex-specific k-mer signals. Genes strongly supported as autosomal by SDpop were classified as not sex-linked (NSL). Genes with low SDpop support for sex linkage (≤ 0.6) that were not strongly supported as autosomal, as well as genes with limited genotype information were classified as inconclusive (Supplementary Data Table S2).

For each gene, the posterior probabilities of autosomal and sex linkage (XY or ZW) inheritance were visualized as scatter plots using ggplot2 in R along each chromosome of the *P. dactylifera* reference genome BC4. Distinct data layers (SDpop autosomal and sex linkage posterior probabilities, iKISS sex-linked kmer -log10, and dS values) were visualized along each chromosome of the *P. dactylifera* BC4 reference genome using Circos plots in shinyCircos v2.0 (Wang Yazhou *et al*. 2023).

Because a reference genome for *P. aequatorialis* is not available, we assessed the conservation of the genomic region of the *P. dactylifera* (BC4) reference genome harboring *P. aequatorialis* sex-linked genes across more distantly related palm species. We elaborated this by constructing synteny maps among three publicly available genomes: *P. dactylifera* (BC4; NCBI BioProject PRJNA83433), *Chamaerops humilis* L. (NCBI BioProject PRJNA106610847), and *Elaeis guineensis* Jacq. (NCBI BioProject PRJNA636092), using MCScan (Python version) (https://github.com/tanghaibao/jcvi/wiki/Mcscan-(python-version)) after extraction of CDS sequences with gffread v0.12.7 (Pertea and Pertea 2020).

### Exon fixed SNPs by sex

For each SNP detected within the captured exons, we quantified the number of male and female individuals that were homozygous and heterozygous for allele 1 and allele 2 of the SNP, based on individual level genotype and allele information provided by the SDpop output. Under an XY sex chromosome system (selected based on the BIC score), sex-linked SNPs are expected to be homozygous in females (XX) and heterozygous in males (XY). Accordingly, a SNP is considered as sex-fixed when all 11 studied females and all 10 studied males were homozygous and heterozygous for the same SNP, respectively. Under a ZW sex chromosome system (selected based on the BIC score), the opposite pattern (heterozygous females and homozygous males) must be considered. We computed for each gene the total number of SNPs and the subset of SNPs showing sex fixed genotypes. The distribution of total and sex fixed SNPs across chromosomes of the *P. dactylifera* BC4 reference genome was visualized as heatmaps using the Python library matplotlib (Hunter 2007).

### Candidate sex-linked markers in *Phytelephas*

We developed SNP based markers from three candidate HCSL genes (Supplementary Data Table S3) to discriminate between male and female *Phytelephas* individuals. In addition, two autosomal loci were included as internal controls to verify PCR performance and DNA quality. Primers were designed to target fixed sex specific SNPs within the HCSL genes, with particular emphasis on positioning the fixed SNPs nucleotide near the 3′ end of the primer to maximize allelic discrimination and amplification specificity. All primer sequences and targeted SNP positions are provided in Supplementary Data Table S3. Genomic DNA was extracted from silica dried leaf tissue of 11 female and 10 male individuals (Supplementary Data Table S1 (VSLG)) sampled from multiple wild populations across Ecuador using the DNeasy Plant Mini Kit (Qiagen, Hilden, Germany), following the manufacturer’s protocol with minor modifications to improve DNA yield and purity. DNA concentration and quality were assessed by nanodrop. PCR amplifications were performed in 40 µL reaction volumes containing approximately 20–50 ng of template DNA, 1× GoTaq® G2 Reaction Buffer (Promega, Madison, WI, USA), 0.2 mM of each dNTP, 0.2 µM of each primer, and 1 U of GoTaq® G2 DNA Polymerase. Thermocycling conditions consisted of an initial denaturation at 95 °C for 2 min, followed by 30 cycles of denaturation at 95 °C for 1 min, annealing at 61 °C for 1 min, and extension at 72 °C for 1 min, with a final extension step at 72 °C for 5 min. PCR products were visualized by electrophoresis on 2% agarose gels.

## Results

### Exon capture data evaluation

Exon capture reads from *Phytelephas aequatorialis* samples were mapped to the reference genome of *Phoenix dactylifera* BC4 to extract SNPs for downstream inference of sex specific genotypes using SDpop. Given the significant phylogenetic divergence between *Phytelephas* and *Phoenix*, we first evaluated the mappability and coverage of the exon capture data by quantifying the number of mapped reads and the number of genes recovered when mapping *Phytelephas* exon capture reads to the *P. dactylifera* reference genome.

Across the 21 samples analyzed (Pop-ExC), the number of reads mapped and properly paired to the *P. dactylifera* reference genome ranged from 19 741 991 to 30 077 767 (mapping rate: 95.3 – 97.8%; standard deviation: 0.78) and from 19 086 912 to 29 066 106 (properly paired rate: 92.2 – 95.3%; standard deviation: 1.14), respectively (Supplementary Data Table S4). The mean number of genes recovered across samples was 29 739, corresponding to approximately 86% (standard deviation: 1.006) (Supplementary Data Table S4) of the predicted genes in the *P. dactylifera* BC4 reference genome.

These results indicate a good conservation of exon sequences between *Phytelephas aequatorialis* and *Phoenix dactylifera*, despite belonging to two different subfamilies within the Arecaceae. This level of cross species exon capture and mapping efficiency supports the reliability of the recovered exon data for calling informative SNPs and provides sufficient genomic representation for robust inference of sex-linked genes.

### Validity of the genetic structure and relatedness of *P. aequatorialis* samples

Before inferring the sexual system and identifying sex-linked genes, we examined how genetically similar the sampled individuals were to each other and whether the samples showed any genetic structure. After filtering variants by read depth, retaining only sites with a minimum coverage of seven reads per individual, a total of 784 686 SNPs and indels were used to assess genetic relatedness among individuals and their genetic structure.

Pairwise relatedness among samples ranged between 0.189 and 0.225 (Supplementary Data Table S5; Supplementary Data Fig. 1A) and did not show any bias regarding the sex of the samples. These values suggest that individuals are neither clones nor duplicated samples and do not include unusually closely related pairs. This level of relatedness is appropriate for downstream analyses of sex linkage, as it minimizes biases in allele frequency based inference.

The genetic structure of the *Phytelephas* studied individuals, as revealed by PCA (Supplementary Data Fig. 1B), showed weak overall genetic differentiation among samples. The first two principal components presented 5.63% (PC1) and 5.51% (PC2) of the total genetic variance, respectively. Most individuals clustered closely together, indicating low genetic differentiation across the majority of the dataset. Two female individuals (Pha13 and Pha16) were clearly separated from the main cluster along PC1 and PC2, suggesting higher genetic differentiation compared to the other samples. However, the relatedness estimates indicated that Pha13 and Pha16 did not show abnormal levels of genetic similarity to other samples compared with the rest of the dataset. Therefore, all individuals were retained for downstream analyses.

### Determination of sex chromosome type, sex-linked genes and sex-linked region

Based on the exon capture data, the sex chromosome system of *P. aequatorialis* was inferred using SDpop by comparing models assuming ZW or XY systems, as well as a model suggesting the absence of sex linkage. According to the Bayesian Information Criterion (BIC), the XY model provided the best fit to the data, as it yielded the lowest BIC value (Supplementary Data Table S6). Approximately, 83% of the polymorphic sites detected in the *Phytelephas* data were inferred to be autosomal, whereas the proportions of sex-linked (XY gametologous) and haploid sites were estimated at about 0.12% and 0.36%, respectively. Paralogous sites accounted for around 1.7% of the total sites.

Sex-linked polymorphisms were investigated using three independent and complementary approaches; SDpop, Synonymous divergence (dS), and iKISS. These methods consistently identified sex-linked genes in *P. aequatorialis*, which were predominantly mapped to a single continuous region on chromosome NC_052409.1 (Chromosome 18) of the *P. dactylifera* BC4 reference genome. This region spans approximately 4.8–5.9 Mb (Figs. 2 and 3). Synteny analysis showed that this region corresponds to homologous continuous regions in other palm species, including *Elaeis guineensis* and *Chamaerops humilis* (Supplementary Data Fig. S2).

**Figure 2:**
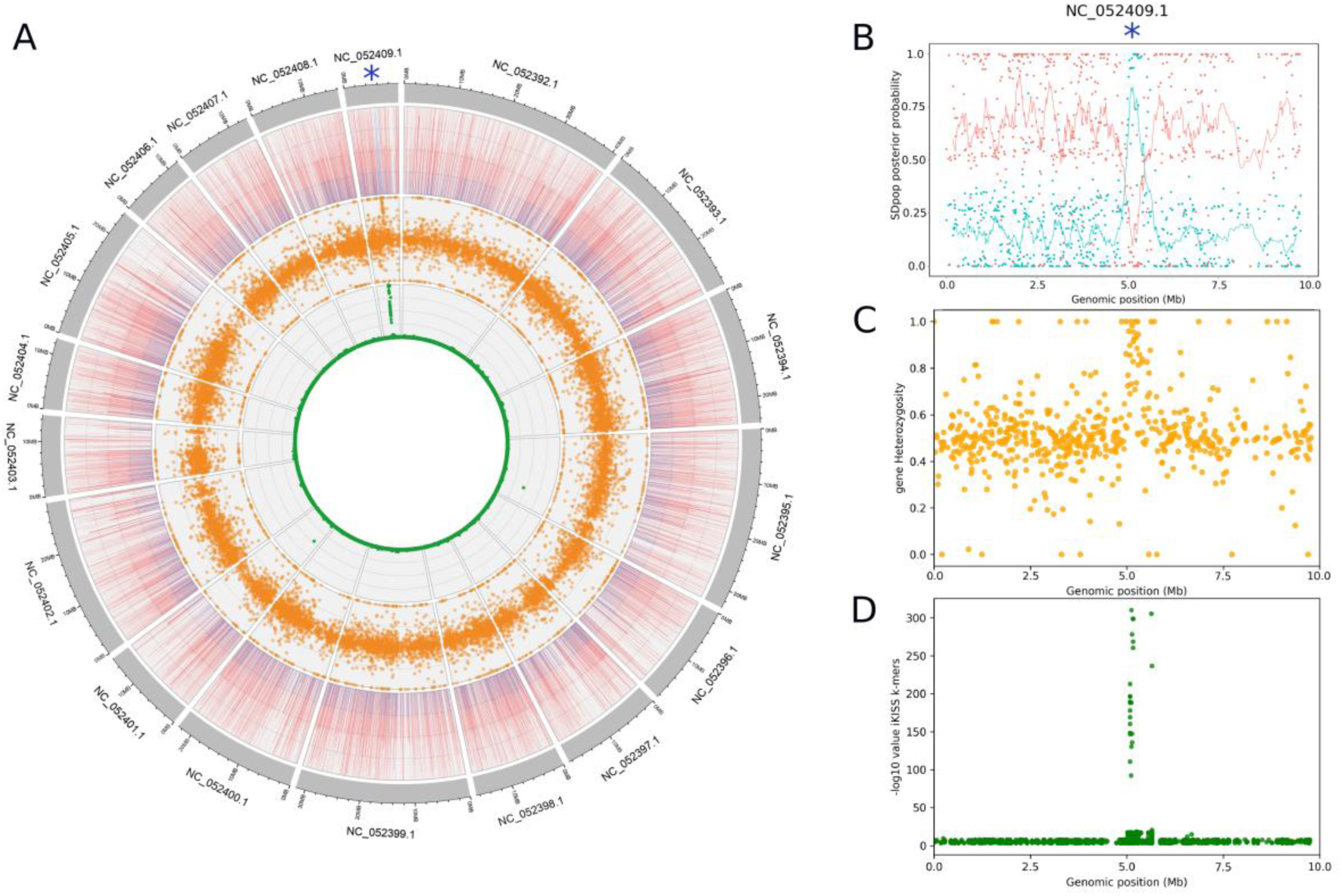
Identification of the sex-linked region in *Phytelephas aequatorialis* mapped onto the reference genome *of Phoenix dactylifera* BC4. **(A)** Circos plot showing, from the outermost to the innermost tracks, the chromosomes of the reference genome, the SDpop posterior probability of genes being autosomal (red bars) or XY sex-linked (blue bars), gene heterozygosity values, and the −log10 k-mer association scores for male or female assignment obtained with iKISS. The blue asterisk indicates the sex-linked region in *Phytelephas* supported by all three methods. **(B)** Zoomed view of the sex-linked region on chromosome 18 (NC_052409.1) of the BC4 genome, showing the SDpop posterior probabilities for genes being autosomal (red bars) or XY sex-linked (blue bars). **(C)** and **(D)** Gene heterozygosity values and −log10 k-mer association scores for male and female assignment inferred by iKISS along chromosome 18 (NC_052409.1) of the reference genome, respectively.

**Figure 3:**
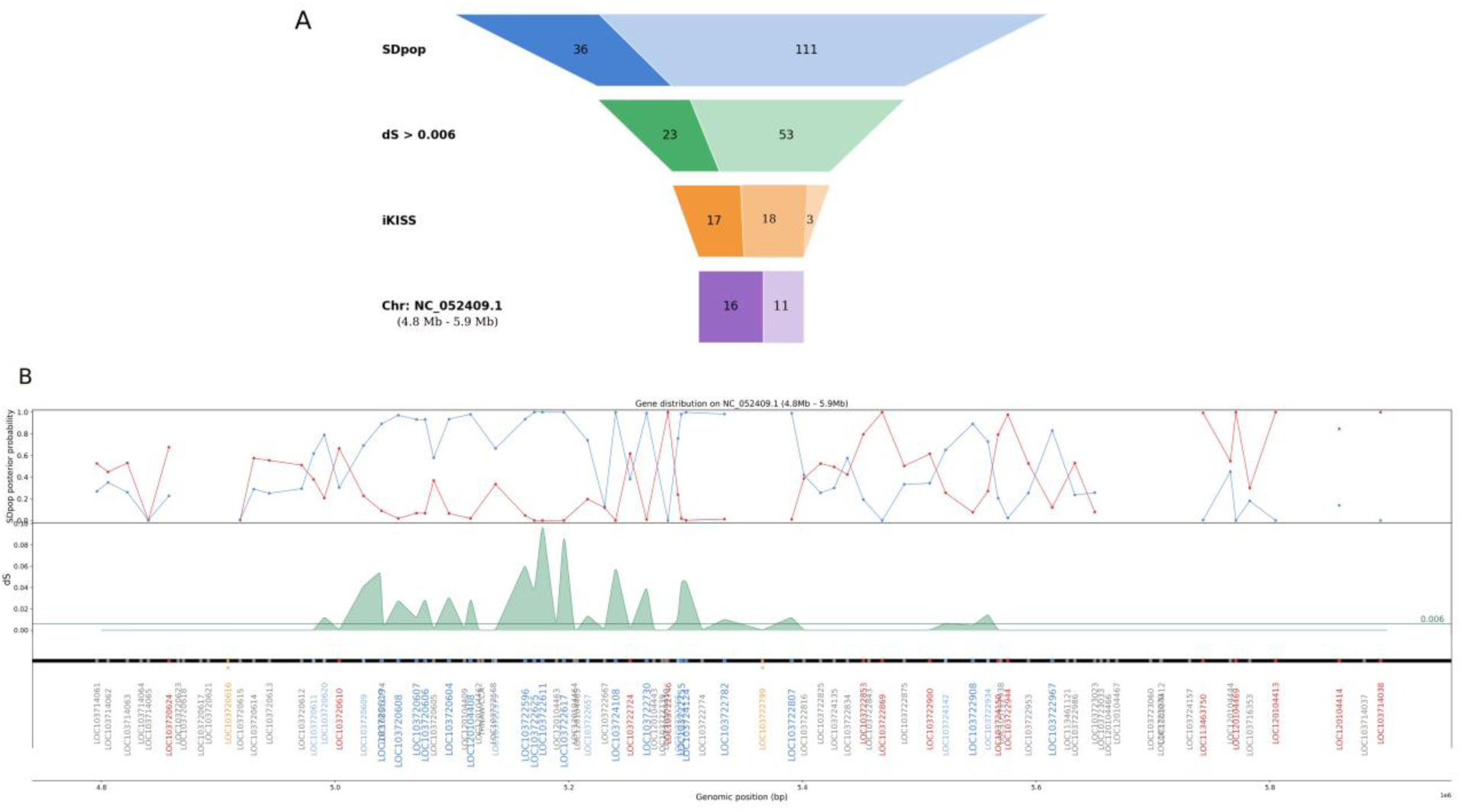
Identification of high confidence sex-linked (HCSL) and putative sex-linked (PSL) genes in *Phytelephas*, with a focus on the sex-linked region spanning 4.8–5.9 Mb on chromosome 18 (NC_052409.1) of *P. dactylifera* BC4 reference genome. **(A)** Funnel plot showing the number of genes retained after each approach used for identifying HCSL and PSL. Dark and light blue indicate genes with SDpop posterior probabilities ≥0.8 and 0.6–0.8, respectively. Dark and light green represent genes meeting the same SDpop thresholds with dS > 0.006. Dark orange indicates genes additionally supported by iKISS sex assignment, whereas less dark orange represents genes supported only by iKISS and light orange for the genes showing SDpop posterior probability 0.6– 0.8, dS>0.006 and assigned a sex by iKISS. Dark purple and light purple indicate HCSL and PSL genes located within the 4.8–5.9 Mb sex-linked region, respectively. See Materials and Methods for HCSL and PSL classification criteria. **(B)** SDpop posterior probabilities for autosomal (red) and XY sex-linked (blue) genes, together with dS values across the 4.8–5.9 Mb region. The horizontal line indicates the genome wide heterozygosity threshold (dS = 0.006). Gene labels follow the reference genome annotation and are color coded as follows: grey, inconclusive; red, autosomal; dark blue, SDpop posterior probability ≥0.8; light blue, SDpop posterior probability 0.6–0.8; orange, genes supported only by iKISS, with no evidence from SDpop or dS.

Based on SDpop inference, 36 genes showed a high posterior probability (≥ 0.8) of being sex-linked, of which 19 mapped to this region (Figs. 2 and 3; Supplementary Data Table S7). The remaining genes with high posterior probability were located on other chromosomes (10 genes) or on unplaced scaffolds (7 genes) in the *P. dactylifera* BC4 reference genome. In addition, 111 genes with intermediate posterior probabilities (> 0.6 and < 0.8) were distributed across multiple chromosomes of the reference genome, *P. dactylifera* BC4 (Fig. 2, Supplementary Data Figs. S2 and S3).

Synonymous divergence (dS) provided additional support for sex linkage among genes identified by SDpop. For genes with SDpop posterior probabilities > 0.6, dS values ranged from 0 to ca. 1.3. A dS threshold of 0.006 was defined based on the median heterozygosity per gene calculated across all genes included in the SDpop analysis (Supplementary Data Table S8). Mapping median heterozygosity per gene onto the *P. dactylifera* (BC4) reference genome revealed a peak in the region identified as sex-linked on chromosome 18 (Fig. 2). In total, 76 genes (XY linkage posterior probabilities > 0.6) exhibited dS values exceeding this threshold (Fig. 3). Among those located in the identified sex-linked region on chromosome 18 of the *P. dactylifera* BC4 reference genome, only two genes showed dS value below the threshold (Fig. 3; Supplementary Data Table S7). Of the 76 candidate genes, 23 exhibited strong evidence of sex linkage (SDpop posterior probability ≥ 0.8), whereas 53 genes showed moderate support for sex linkage, with posterior probabilities between 0.6 and 0.8 (Fig. 3; Supplementary Data Table S7).

Independent k-mer based analysis using iKISS further supported the localization of sex-linked sequences to the same genomic region on chromosome NC_052409.1 (chromosome 18) of the reference genome *P. dactylifera* (BC4) (Figs. 2 and 3). Sex-linked contigs assembled from *Phytelephas* k-mers and detected by iKISS to be sex-linked based on the highest −log(p) values were predominantly concentrated within the 4.8–5.9 Mb interval of NC_052409.1 (Figs. 2 and 3). After BLAST mapping of iKISS sex-linked contigs to the *P. dactylifera* BC4 reference genome, contigs were assigned to genes (Supplementary Data Table S9). Some contigs matched genes inferred by SDpop to be autosomal (posterior probability > 0.8), likely reflecting spurious alignments due to the short length of the assembled contigs by iKISS (100–312 bp). Because these contigs showed conflicting assignments (identified as sex-linked by iKISS but mapping to genes inferred to be autosomal by SDpop), they were considered ambiguous and excluded from subsequent analyses. After filtering, 38 genes were supported as sex-linked by iKISS. Among these, 17 genes showed strong evidence of sex linkage, with an SDpop posterior probability ≥ 0.8 and dS value > 0.006. The remaining 21 genes were supported either by iKISS alone (18 genes) or by moderate evidence from SDpop (posterior probability between 0.6 and 0.8) and dS value > 0.006 (3 genes) (Fig. 3; Supplementary Data Table S7). Of the 38 genes, 22 genes had mapped to chromosome NC_052409.1 of the *Phoenix* reference genome (BC4) within the 4.8–5.9 Mb interval. The remaining genes were located on other chromosomes or on unplaced scaffolds.

By integrating evidence from SDpop, dS, and iKISS, we identified 17 genes with high confidence as sex-linked (HCSL) (classification criteria in Supplementary Data Table S2). Genes were classified as HCSL when they were supported by SDpop (XY posterior probability ≥ 0.8), dS > 0.006, and iKISS (sex-linked contigs after filtering of ambiguous and autosomal assignments). Sixteen of these genes clustered within a single genomic region of approximately 365 kb within the previously identified sex-linked region (positions 5 030 459 – 5 395 612, Figs. 2 and 3) on chromosome NC_052409.1 (chromosome 18) of the *P. dactylifera* BC4 reference genome, while the only remaining gene was located on an unplaced scaffold (Table 1). In addition, 11 potentially sex-linked genes (PSL) were detected on the identified sex-linked region of the chromosome 18 based on less stringent criteria (classification criteria in Supplementary Data Table S2); Supplementary Data Table S7; Fig. 3).

**Table 1:** High-confidence sex-linked (HCSL) genes in *Phytelephas aequatorialis*, their syntenic positions in the *Phoenix dactylifera* reference genome (BC4), and functional annotations. (gene IDs were inferred from the reference genome annotation of *P. dactylifera* (BC4))

| gene | chromosome | start | end | XY_prior | noprior_autosomal | dS | ikiss_Tag | Function | Class |
| --- | --- | --- | --- | --- | --- | --- | --- | --- | --- |
| LOC103720619 | NC_052409.1 | 5030459 | 5048874 | 0.893104 | 0.0897395 | 0.056 | MALE | histidine kinase 1, transcript variant X2 | HCSL |
| LOC103720608 | NC_052409.1 | 5051405 | 5056542 | 0.971909 | 0.0184431 | 0.0283 | MALE | proteasome subunit alpha type-7 | HCSL |
| LOC103720607 | NC_052409.1 | 5068967 | 5070178 | 0.932205 | 0.0677909 | 0.0113 | MALE | myb-related protein Hv1 | HCSL |
| LOC103720606 | NC_052409.1 | 5075953 | 5077984 | 0.932487 | 0.0675096 | 0.0302 | MALE | protein POLLENLESS 3-LIKE 2 | HCSL |
| LOC103720604 | NC_052409.1 | 5089132 | 5105562 | 0.935593 | 0.0643869 | 0.0321 | MALE | synaptotagmin-2-like | HCSL |
| LOC120104408 | NC_052409.1 | 5112993 | 5118655 | 0.980546 | 0.0189312 | 0.032 | MALE | 3-dehydroquinate synthase, chloroplastic-like, transcript variant X2 | HCSL |
| LOC103722596 | NC_052409.1 | 5158941 | 5166166 | 0.935216 | 0.0461037 | 0.0618 | MALE | pre-mRNA-processing factor 17-like, transcript variant X1 | HCSL |
| LOC103722625 | NC_052409.1 | 5167807 | 5173012 | 0.999994 | 6.43E-06 | 0.0345 | MALE | tRNA dimethylallyltransferase 9, transcript variant X2 | HCSL |
| LOC103722611 | NC_052409.1 | 5174376 | 5180426 | 0.999999 | 9.56E-07 | 0.1027 | MALE | SKP1-like protein 1A | HCSL |
| LOC103722617 | NC_052409.1 | 5192774 | 5198696 | 0.999497 | 0.000367019 | 0.0945 | MALE | SKP1-like protein 1 | HCSL |
| LOC103724108 | NC_052409.1 | 5239347 | 5240602 | 0.999717 | 0.000282364 | 0.0609 | MALE | bZIP transcription factor 27 | HCSL |
| LOC103722730 | NC_052409.1 | 5264996 | 5268065 | 0.991489 | 0.00825589 | 0.0418 | MALE | transcription factor ABORTED MICROSPORES, transcript variant X2 | HCSL |
| LOC103722755 | NC_052409.1 | 5295670 | 5296609 | 0.981614 | 0.0177305 | 0.0467 | MALE | uncharacterized LOC103722755 | HCSL |
| LOC103724124 | NC_052409.1 | 5296772 | 5303749 | 0.998691 | 0.00130862 | 0.0464 | MALE | uncharacterized LOC103724124, transcript variant X2 | HCSL |
| LOC103722782 | NC_052409.1 | 5322204 | 5344121 | 0.982578 | 0.0124966 | 0.0101 | MALE | ribonuclease P protein subunit p25-like protein | HCSL |
| LOC103722807 | NC_052409.1 | 5385661 | 5395612 | 0.99036 | 0.00962884 | 0.0122 | MALE | probable protein phosphatase 2C 60 | HCSL |
| LOC103719241 | NW_024067747.1 | 60147 | 62611 | 0.993425 | 0.00500507 | 0.0935 | MALE | 1-Cys peroxiredoxin-like | HCSL |

The HCSL genes encompassed several functional categories, including transcriptional regulators (e.g. MYB-related protein Hv1 and bZIP transcription factor 27), components of protein turnover pathways (e.g. proteasome subunit alpha type-7 and S-phase kinase-associated protein 1 (SKP1-like) proteins), and signaling-related proteins (e.g. histidine kinase 1 and protein phosphatase 2C). Additional genes were associated with RNA processing and translation (e.g. pre-mRNA-processing factor 17-like, ribonuclease P subunit p25-like and tRNA dimethylallyltransferase 9), cellular trafficking (e.g. synaptotagmin-2-like), and metabolic or redox functions (e.g. 3-dehydroquinate synthase and 1-Cys peroxiredoxin) (Table 1).

### Development of sex markers for early sex detection in *Phytelephas*

We designed five primer pairs, three of which were designed from the Y haplotype sequences of *P. aequatorialis* identified by SDpop to differentiate male and female samples, whereas the remaining two primer pairs target autosomal regions and therefore serve as PCR controls. The three genes (LOC103722625 (Dimet); LOC103720606 (POLY); LOC103722617 (SKP)) selected were among those previously assigned as HCSL (Table 1, Supplementary Data Table S3, gene IDs were inferred from the reference genome annotation of *P. dactylifera* (BC4)). Primer locations were based on male sex-fixed SNPs identified in these genes from the DNA capture data. A SNP was considered male sex-fixed when all analyzed male individuals carried the same male-specific allele, which was absent from all analyzed female individuals. Among the 16 HCSL genes, 15 contained sex fixed SNPs (Supplementary Data Table S10, Supplementary Data Fig. S4).

The developed sex markers targeting the male allele of the Y haplotype were validated on 21 individuals from eight populations (Fig. 1, Supplementary Data Table S1(VSLG)) by PCR and revealed by agarose gel electrophoreses. They showed a DNA amplification in all male individuals, whereas no amplification was detected in any female individuals thereby providing a reliable tool for sex discrimination in the species (details on each marker were reported in Supplementary Data Table S3 and amplification results on gel electrophoreses are illustrated in Supplementary Data Fig. S5).

These results provide further support for the sex-linked nature of these genes, as the associated markers were validated in an independent set of *P. aequatorialis* individuals not included in the initial identification of the sex-linked regions

## Discussion

### Identification of an XY sex chromosome system in the ivory palm

Based on exon capture data from *P. aequatorialis* individuals, we found evidence consistent with an XY sex chromosome system in this species, as indicated by the BIC values estimated by SDpop. Despite that several palm species are dioecious (30% of dioecious species in the family (Nadot *et al*. 2016)), their sex chromosome systems remain poorly studied. Within the family, the sex chromosomes of *Phoenix* dactylifera have received most attention, and a XY system has been identified and shown to be conserved across all the genus *Phoenix* (Cherif *et al*. 2016; Renner 2014; Torres *et al*. 2018; Tessarotto *et al*. 2025). Recently, a XY sex chromosome system was identified in *Kerriodoxa* (Tessarotto *et al*. 2025), a genus belonging to the same subfamily as *Phoenix* (subfamily Coryphoideae) and their common recent ancestor was dated to approximately 66 million years ago (My) (Nadot *et al*. 2016). Here we report the first evidence of an XY sex chromosome system in the subfamily Ceroxyloideae, which is predominantly composed of dioecious species and diverged from Coryphoideae approximately 87 My. Therefore, in all the three dioecious palm species for which sex chromosomes have been identified to date, the sex chromosome system is XY, as is the case for the majority of dioecious plant species (ca. 80-85% of the cases) for which sex chromosomes are characterized (Renner and Müller 2022; Lesaffre *et al*. 2024).

The evolution of dioecy and sex chromosomes has traditionally been explained by the “two gene model” proposed by Charlesworth & Charlesworth (1978), in which linked mutations causing male and female sterility, followed by recombination suppression, give rise to differentiated sex chromosomes (Charlesworth 2016). However, recent evidence suggests that this model may not be universal in plants, as several species appear to possess small sex-determining regions controlled by a single master regulator rather than two linked sterility mutations (Renner and Müller 2021). These findings highlight the diversity of evolutionary pathways leading to dioecy and underscore the need to investigate the genetic architecture of sex determination in each lineage including palms.

### A new genomic sex-linked region *in P. aequatorialis* distinct from known sex-linked regions in palms

As a reference genome for *P. aequatorialis* is currently unavailable and we aimed to compare sex-linked regions across palm species, following Tessarotto et al. (2025), we mapped the *P. aequatorialis* sequences to the *P. dactylifera* BC4 reference genome.

Using exon capture data of *P. aequatorialis*, we applied three independent and complementary approaches (SDpop, dS analysis, and iKISS) to identify sex-linked genes. Convergently, the three approaches identified 17 HCSL genes and 90 PSL genes, the latter supported by one or two of the methods (genes classification details in Supplementary Data Table S2). The consistency of the results across these complementary approaches strengthens the inference that these genes are associated with sex linkage. In addition, ca. 94% of the HCSL genes clustered within a single genomic region spanning approximately 365 kb on chromosome 18 (NC_052409.1) of the *P. dactylifera* (BC4) reference genome, supporting its identification as a very promising candidate sex-linked region. In contrast, only ca. 12% of PSL genes (11 over 90 genes) were located in this region. The presence of fixed sex-linked SNPs across all HCSL genes provides additional evidence that these loci are closely associated with sex determination or located within a non-recombining genomic region linked to the sex determining locus (Supplementary Data Fig. S4, Supplementary Data Table S10). Importantly, comparative analyses indicate that the genomic region identified here as sex-linked is conserved across the two palm species, *Elaeis guineensis* (GenBank accession: GCA_015461965.1) and *Chamaerops humilis* (GenBank accession: GCA_042465325.1), suggesting that synteny is maintained in this portion of the genome across the family. The conservation of gene order across these phylogenetically distant palms supports the interpretation that this genomic segment is evolutionarily stable and therefore would possibly be conserved in *P. aequatorialis* genome as well.

In *Kerriodoxa*, the sex-linked region corresponded to the same genomic region identified in *Phoenix* on the chromosome 12 of the reference genome *P. dactylifera* BC4 (Tessarotto *et al*. 2025). This discovery raises the possibility of convergent evolution of this region as sex-linked between these two species (Tessarotto *et al*. 2025). In contrast, the candidate sex-liked region identified in *P. aequatorialis* maps to chromosome 18 of the *P. dactylifera* genome (BC4), suggesting that different genomic regions may have been recruited as sex determining loci in different palm subfamilies.

A dynamic evolution of sex chromosome and sex determining regions has been well documented in flowering plants. For example, in the genus *Populus*, the sex determining region is located on chromosome 19 in *P. trichocarpa* and *P. tremula*, whereas it occurs on chromosome 14 in *P. euphratica* (Müller *et al*. 2020; Yang *et al*. 2021). Similarly, in the genus *Salix*, different species possess sex determining regions on distinct chromosomes, such as chromosomes 7 and 15 in *S. chaenomeloides* and *S. arbutifolia*, respectively (Wang *et al*. 2022; Wang Yi *et al*. 2023; Yang *et al*. 2021). It would be of particular interest to investigate whether the same genomic region underlies sex chromosomes throughout the predominantly dioecious Ceroxyloideae palm subfamily, and whether dioecy in these species originated from a common dioecious ancestor.

The congruent results obtained from the complementary approaches used here (SDpop, dS, and iKISS) provided robust evidence for the existence of a sex-linked region in *Phytelephas*, consistent with an XY system, and allowed us to identify candidate sex-linked genes with confidence. This level of resolution was sufficient for an initial characterization of the sex-linked region and sex-linked genes in this species. However, to further characterize the structure and evolution of the sex-linked region, including potential structural rearrangements between X and Y haplotypes, repetitive element accumulation, and epigenetic modifications, we will assemble chromosome-level reference genomes from male and female individuals of *Phytelephas*.

### Candidate sex-linked genes in *P. aequatorialis*

The putative functions of the 17 HCSL and 90 PSL genes identified using the three complementary approaches were inferred from the reference genome annotation of *P. dactylifera* (BC4). Numerous HCSL genes identified in *P. aequatorialis* have functions associated with reproductive development. In particular, we identified homologs of ABORTED MICROSPORES, a transcription factor involved in microspore development and pollen formation, as well as homologs of POLLENLESS-like proteins implicated in male fertility and pollen development in plants (Glover *et al*. 1998; Sorensen *et al*. 2003; Xu *et al*. 2010). Genes involved in pollen and gametophyte development have been described as implicated in the evolution of dioecy, as mutations affecting male or female fertility can initiate the separation of sexes and subsequently become linked to sex determining regions (Shi *et al*. 2015; Charlesworth 2016). Similar patterns have been observed in other dioecious palms. In *P. dactylifera*, the Y chromosome contains a male specific region harboring genes required for male reproductive development, including Cytochrome P450 703 (CYP703), involved in sporopollenin synthesis and pollen wall formation, and Glycerol-3-phosphate acyltransferase 3 (GPAT3), which plays a key role in anther and pollen development (Torres *et al*. 2018; Leite Montalvão *et al*. 2021).

Among the HCSL genes identified in *P. aequatorialis*, the presence of two SKP1-like genes within the sex-linked region is particularly noteworthy. SKP1 proteins are core components of the SCF (SKP1–Cullin–F-box) ubiquitin ligase complex, which mediates targeted protein degradation and regulates numerous developmental processes in plants. In *Arabidopsis*, the SKP1 homolog ASK1 is essential for male meiosis and pollen development, and disruption of this gene results in male sterility (Yang *et al*. 1999; Zhao *et al*. 2003). The occurrence of SKP1-like homologs showing to be HCSL and present within the sex-linked region identified for *P. aequatorialis* may therefore represent an important regulatory component of this region.

Additional HCSL candidates include transcriptional regulators such as MYB-related and bZIP transcription factors, which regulate diverse developmental processes including floral organ differentiation and reproductive development (Chopy *et al*. 2023; Wu *et al*. 2024). Signaling components such as histidine kinase 1, a component of cytokinin signaling pathways, were also detected and may contribute to regulatory networks controlling floral development and sex expression (Henning 2025). In addition to the HCSL genes, several PSL genes identified in our analysis belong to functional categories frequently associated with reproductive development and regulatory pathways. Notably, a homolog of LONELY GUY (LOG4), a gene involved in cytokinin activation. Cytokinin signaling plays an important role in floral meristem activity and reproductive development, and a LOG gene has also been identified in the sex chromosome of *P. dactylifera*, where it has been proposed to contribute to female flower suppression during the evolution of dioecy (Torres *et al*. 2018; Leite Montalvão *et al*. 2021; Tessarotto *et al*. 2025). Comparative studies in the dioecious palm *Kerriodoxa elegans* further support the involvement of similar functional pathways in sex-linked regions (Tessarotto *et al*. 2025). Although the specific genes associated with sex-linked regions differ among palm genera, these studies collectively indicate that sex-linked regions in palms frequently contain genes involved in pollen development, transcriptional regulation, and key developmental pathways. This pattern is consistent with findings across dioecious plant species, where sex-linked regions are often enriched for genes controlling reproductive development and floral organ differentiation (Muyle *et al*. 2017; Leite Montalvão *et al*. 2021).

Future analyses focusing on the genomic context of these candidate genes, including the characterization of nearby repetitive elements, DNA methylation patterns, and sex biased gene expression in male and female flowers, will provide further insights into the molecular mechanisms underlying sex determination in ivory palms.

### Early sex distinction by SNP markers of ivory palm

Dioecious palms are traditionally distinguished as male or female based on the morphology of their flowers and inflorescences. However, this approach is only possible once individuals reach the stage of reproductive maturity. In palms, the age of first flowering period varies considerably between species, ranging from ca. 4 to 7 years in some specie like the date palm (*P. dactylifera*) (Jani *et al*. 2025) to several decades for example in *Arenga* (ca. 15 years) and *Corypha* (ca. 45 years) (Pongsattayapipat and Barfod 2009; Barfod *et al*. 2011). In *Phytelephas*, demographic studies estimate that reproduction maturity began at about 24 years of the plant age under natural conditions (Bernal 1998). Such long juvenile period limits a morphological sex identification at young stage and represent a major constraint for breeding and management programs.

Several dioecious palms have major ecological and economic importance (Dransfield *et al*. 2008). Among these species we cite for instance, the date palm is one of the most important perennial fruit crops in arid regions, providing a staple food source and supporting agroecosystems across North Africa and the Middle East (Al-Karmadi and Okoh 2024). Because only female individuals produce fruits in dioecious species, early sex identification is highly desirable to optimize cultivation and maximize yield, typically by maintaining a higher proportion of female plants while preserving sufficient males for pollination. For instance, SNP-based sex markers identified by PCR have been developed to distinguish male and female *P. dactylifera* at early developmental stages (Cherif *et al*. 2013). These markers contribute to the efficient management of palm plantations propagated through sexual reproduction.

*Phytelephas aequatorialis* is an ecologically and economically important palm native to northwestern South America, particularly Ecuador (Koziol and Pedersen 1993; Brokamp 2015; Escobar *et al*. 2021). The species contributes to forest ecosystem functioning by providing food resources and habitat for numerous animal species, including seed dispersers and pollinators. In addition to its ecological role, *P. aequatorialis* produces large seeds whose hardened endosperm forms a dense ivory-like material known as vegetable ivory or “tagua”. This material has historically been used in the manufacture of buttons, jewelry, carvings, and handicrafts, and remains a sustainable plant-based alternative to animal ivory, while also showing potential for biotechnological applications (Carvajal-Barriga *et al*. 2022, 2023). The leaves of *Phytelephas* species are also widely used for traditional roof thatching and construction materials in rural communities (Koziol and Pedersen 1993; Pülschen 2000; Escobar *et al*. 2022; Montúfar *et al*. 2022).

These ecological and socio-economic values highlight the importance of improving management and cultivation strategies for this species.

Given the long juvenile phase of *P. aequatorialis*, the development of reliable molecular tools for early sex determination is particularly valuable (Acosta-Solis 1948). Molecular markers based on SNPs distinguishing X and Y haplotypes in sex-linked genomic regions provide a robust framework for this purpose. When combined with simple molecular techniques such as PCR amplification followed by agarose gel electrophoresis, such markers represent a cost-effective and accessible diagnostic method for early-stage sex identification. In this study, we developed three sex-linked molecular markers generating PCR fragments of distinct sizes amplified only in male *P. aequatorialis* plants, allowing a clear discrimination between male and female individuals. These markers could significantly improve planting strategies, breeding programs, and germplasm management for *Phytelephas* by enabling the early selection of female individuals for fruit production and supporting the integration of conservation strategies into agroforestry systems. Future work involving larger population validation and comparative genomic analyses across Ceroxyloideae may further refine these markers and provide insights into the evolution of sex chromosomes in the family Arecaceae.

## Conclusion

Following the previous discovery of possible convergent evolution of the sex-linked region in the two Coryphoideae palm species, *P. dactylifera* and *K. elegans*, we show here that in the Ceroxyloideae species *P. aequatorialis*, a different genomic region has been recruited as the sex-linked region. Within this region, 16 genes were classified as high confidence sex-linked genes, including genes involved in pollen development, transcriptional regulation, and key developmental pathways. In addition, we developed three SNP-based sex markers that enable early sex identification in *Phytelephas* plants. These markers have the potential to improve the management of *Phytelephas* seed production.

As the majority of species in the subfamily Ceroxyloideae are dioecious, we further aim to investigate the sex-linked regions and genes in these palm species to determine whether they share or not homologous sex-linked genomic regions, thereby providing insights into the emergence and evolution of dioecy within this subfamily.

Further comparative genomic analyses of dioecious palm species across the family are needed to better understand the pathways of sex determination and sex chromosome evolution. In addition, future studies are necessary to identify the master genes for sex determination in *Phytelephas* and other dioecious species in the family.

## Supplementary Data

Supplementary data consisting of five figures and ten tables are available in the Zenodo repository at: https://doi.org/10.5281/zenodo.22685484

## Acknowledgement

The authors acknowledge the ISO 9001 certified IRD i-Trop HPC (South Green Platform) at IRD Montpellier for providing HPC resources that have contributed to the research results reported within this paper.

The authors thank Mrs Amelie Portalez and Mrs Lynn Wheibe for their contributions to the laboratory work for validating the developed markers by PCR. We want to thank ARCAD (Agropolis Resource Centre for Crop Conservation, Adaptation and Diversity) for elaborating the exon-capture laboratory work, specially we are grateful to the late Mr. Sylvain Santoni for his invaluable contribution to this work. We thank the Otonga Foundation for providing access to the forest reserve and Mr. Arturo Guasti for his assistance in the field.

We would like to thank the herbarium QCA of Pontificia Universidad Católica del Ecuador for providing part of Phytelephas samples used in this study. This research was conducted under the Genetic Resources Access Framework Agreement No. MAATE-DBI-CM-2023-0292 and MAE-DNB-CM-2018-0082, issued by the Ministry of Environment, Water and Ecological Transition of Ecuador.

## AI assistance

ChatGPT (OpenAI) was used to assist with English-language editing of the manuscript.

## Data availability

Exon captures raw data is available on the NCBI.

## AUTHOR CONTRIBUTIONS

F.A. conceived the project and acquired funding. R.M., R.G. and G.O-V. sampled the sequenced palms. T.B. supervised the sample preparation and sequencing. P.F, J.O, H.T and J.K developed the bioinformatic pipelines. P.F. and J.O. performed the analyses of the genomic sequences. P.F. wrote the manuscript. All authors read and approved of the final version of the manuscript.

